# Activated protein C promotes β-arrestin-2- and c-Src-dependent caveolin-1 (Cav1) phosphorylation and alters Cav1 association with PAR1 and GRK5

**DOI:** 10.1101/2025.11.02.686166

**Authors:** Huaping Qin, Lennis B. Orduña-Castillo, Olivia Molinar-Inglis, Monica L. Gonzalez Ramirez, Miguel A. Lopez-Ramirez, Carolyne Bardeleben, JoAnn Trejo

**Affiliations:** Department of Pharmacology and; Department of Medicine, School of Medicine, University of California, San Diego, La Jolla, CA 92093

**Keywords:** APC, arrestin, biased signaling, Cav1, GPCR, GPCR kinase, GRK, protease-activated receptor-1, c-Src, thrombin

## Abstract

G protein-coupled receptors (GPCRs) display bias towards either G proteins or GPCR kinase (GRK)-mediated β-arrestin signaling depending on the agonist stabilized receptor conformation and cellular context. The cellular location of GPCRs particularly within plasma membrane microdomains such as lipid rafts contributes to biased signaling through mechanisms that are not well understood. The protease-activated receptor-1 (PAR1) exhibits biased signaling in response to thrombin and activated protein C (APC). APC-induced β-arrestin-2 (βarr2) biased signaling requires PAR1 compartmentalization in caveolae, a subtype of lipid rafts, whereas thrombin-activated PAR1 G protein signaling does not. Caveolin-1 (Cav1) is the principal structural protein of caveolae and regulates signaling through protein-protein interactions. The mechanisms by which Cav1 contributes to APC/PAR1-induced βarr2 biased signaling is not known. Here we report that APC-activated PAR1 modulates Cav1 phosphorylation via a βarr2- and c-Src-dependent pathway. APC also regulates the dynamics of endogenous PAR1-Cav1 and GRK5-Cav1 co-localization examined by single molecule super-resolution stochastic optical reconstruction microscopy imaging in human cultured endothelial cells. We further demonstrate that GRK5 interacts with Cav1 in intact cells through an N-terminus aromatic-rich consensus Cav1 binding motif. Unlike wildtype GRK5, a GRK5 mutant defective in Cav1 binding localized predominantly to the cytoplasm rather than the plasma membrane and failed to promote βarr2 recruitment to APC-activated PAR1. These studies suggest that Cav1 itself contributes to the regulation of APC-activated PAR1 βarr2 biased signaling likely through multiple mechanisms that may converge on GRK5.

## Introduction

Protease-activated receptor-1 (PAR1) is a G protein-coupled receptor (GPCR) that is activated by the coagulant protease thrombin (1) or the anti-coagulant protease activated protein C (APC) (2) and displays biased signaling (3–6). In endothelial cells, thrombin-activation of PAR1 promotes coupling to heterotrimeric G proteins, leading to barrier disruption and inflammatory responses including upregulation of adhesion molecules and expression of cytokines (7–9). In contrast, APC-activation of PAR1 drives β-arrestin-2 (βarr2)-mediated endothelial cytoprotective responses including barrier stabilization, anti-inflammatory, anti-apoptotic pro-survival activities (3, 10, 11) and βarr2-dependent neuroprotection in a mouse model (12), making recombinant APC a promising drug. Thrombin and APC biased agonism at PAR1 is enabled through distinct cleavage sites of the N-terminus resulting in the generation of unique N-terminus tails. Thrombin binds to and cleaves PAR1 at an N-terminal arginine (R)41 site, generating an activating tethered ligand that activates G protein signaling (13). APC binds to the endothelial protein C receptor (EPCR), a transmembrane cofactor, and cleaves PAR1 at an N-terminal R46 site and activates βarr2 cytoprotective signaling (3, 4, 10, 14).

Caveolae are a subtype of lipid raft plasma membrane microdomains highly abundant in endothelial cells. Previous studies showed that disruption of lipid rafts perturbs localization of PAR1 and EPCR, the APC-cofactor, and blocks APC/PAR1 promoted barrier protection (15, 16). Caveolin-1 (Cav1) is the major structural protein required for caveolae formation and mediates protein-protein interactions to regulate signaling and other cellular functions (17, 18). In addition to lipid raft disruption, we showed that loss of Cav1 expression blocks APC-activated PAR1 endothelial cytoprotective signaling, whereas thrombin-induced PAR1 signaling remains intact (10, 16). These findings indicate that thrombin-vs. APC-activated PAR1 biased signaling occur in distinct plasma membrane microdomains. The mechanism by which distinct plasma membrane microdomains with different lipid composition and enriched proteins like Cav1 regulate GPCR biased signaling is not well defined.

GPCR kinases (GRKs) must localize to the plasma membrane to bind and phosphorylate activated GPCRs and are critical for regulating GPCR biased signaling (19, 20). GRKs use different mechanisms for membrane localization. GRK2 and 3 contain a pleckstrin homology (PH) domain that binds phospholipids and free G protein βγ subunits (21, 22), whereas GRK4, 5 and 6 rely primarily on a C-terminal amphipathic helix (23–25). Besides GPCRs, GRKs are also known to interact with other proteins including Cav1 (26). All GRKs, including GRK5 harbor an N-terminus consensus Cav1 binding motif and GRK2 and GRK5 were shown to interact directly with the Cav1 scaffolding domain (CSD) *in vitro* (27). Moreover, CSD peptides were shown to inhibit GRK2 and GRK5 activity *in vitro* (27). Unlike most biased agonists that rely on different GRKs, we recently showed Th- and APC-activated PAR1 biased signaling use the same GRK, GRK5 (28), despite PAR1 being localized to distinct plasma membrane compartments (i.e. plasma membrane vs. caveolae). Currently, it is not known if Cav1 regulates APC-activated PAR1 βarr2 biased signaling and how this might integrate with GRK5 function.

In this study we found that APC promotes Cav1 phosphorylation through a βarr2- and c-Src-mediated pathway. APC also modulates endogenous PAR1-Cav1 and GRK5-Cav1 colocalization assessed by single molecule super-resolution stochastic optical reconstruction microscopy (STORM) imaging in human cultured endothelial cells. We further demonstrate that GRK5 interacts with Cav1 in intact cells through the previously reported N-terminus consensus Cav1 binding motif (27). A GRK5 mutant defective in Cav1 binding localized predominantly to the cytoplasm rather than the plasma membrane like wildtype GRK5 and failed to promote βarr2 recruitment to APC-activated PAR1. Thus, beyond Cav1’s structural role, these studies suggests that Cav1 interacts with PAR1 and GRK5, key effectors essential for driving APC-activated PAR1 βarr2 biased signaling.

## Results

### APC-activated PAR1 promotes Cav1 Y14 phosphorylation via a c-Src dependent pathway

Cav1 is the principle structural component essential for caveolae formation in endothelial cells. We previously reported that Cav1 expression is required for APC-activated PAR1-βarr2 mediated cytoprotective signaling and not for thrombin-activated PAR1-G protein inflammatory signaling in endothelial cells (10, 16). These studies indicate that activated PAR1 biased signaling is regulated through compartmentalized in distinct plasma membrane microdomains (Fig. 1, *A*). To begin to understand how Cav1 might contribute to APC-activated PAR1 biased signaling we examined Cav1 phosphorylation. Cav1 tyrosine (Y)14 phosphorylation is a key regulatory site that modulates protein-protein interactions (17, 29). To determine if APC-activation of PAR1 modulates Cav1 Y14 phosphorylation, human umbilical vein endothelial cell (HUVEC) derived EA.hy926 cells were stimulated with APC for various times and Cav1 Y14 phoshorylation was assessed by immunoblot. APC stimulated a significant increase in Cav1 Y14 phosphorylation at 30 min that remained elevated for 90 min compared to untreated control cells (Fig. 1, *B* lanes 1-4 and *C*). The kinetics of APC-activated PAR1-βarr2 promoted cytoprotective signaling onset is slow and protracted (3, 10) and is consistent with the timing of APC-induced Cav1 Y14 phosphorylation. These results suggest that APC may modulate Cav1 function.

**Figure 1.**
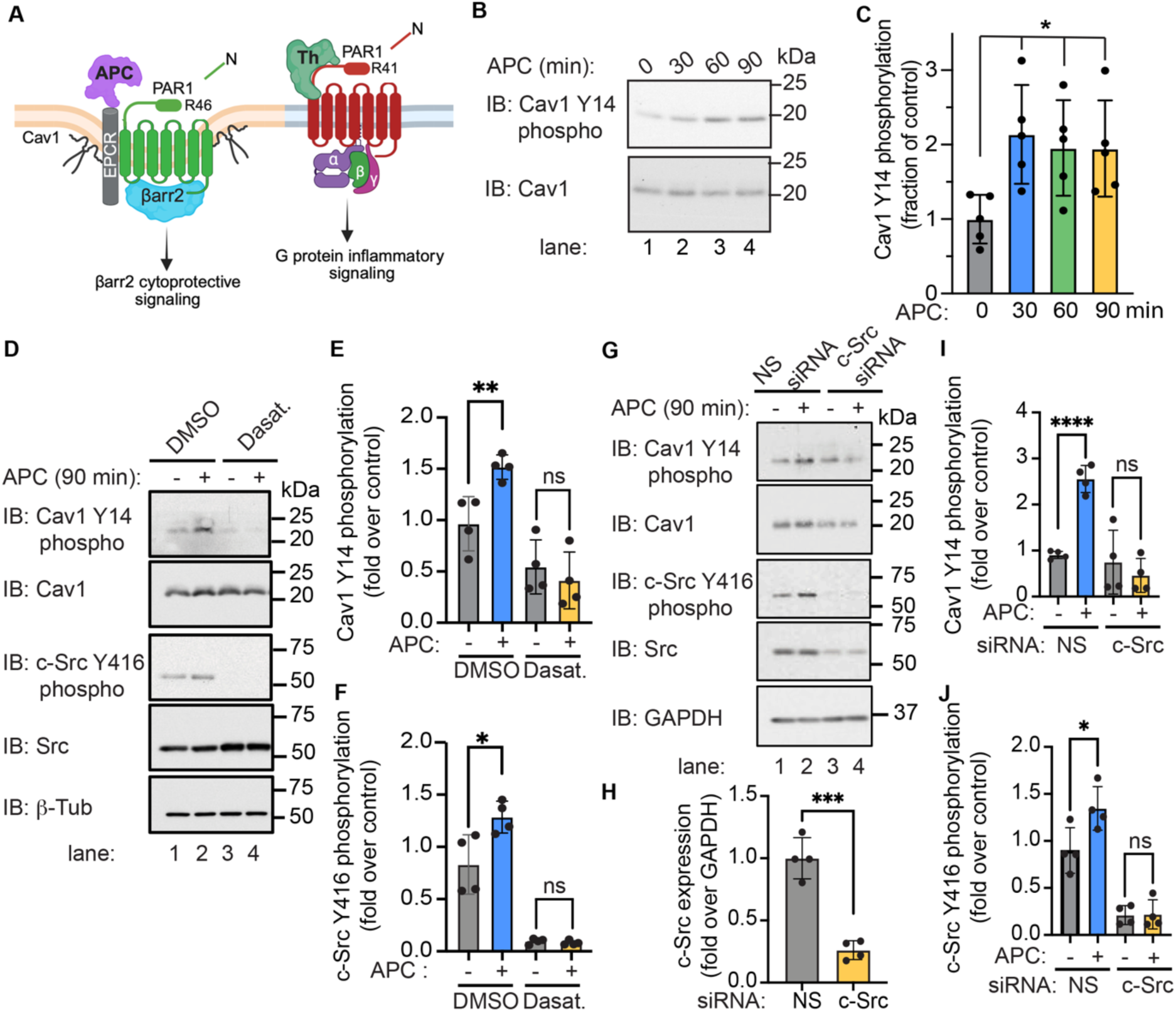
APC-induced Cav1 Y14 phosphorylation is mediated by c-Src. *A*, Cartoon, APC bound to the endothelial protein C receptor (EPCR) cofactor activates PAR1 through arginine (R)46 cleavage in caveolae microdomains and promotes βarr2-driven cytoprotective signaling. Thrombin (Th) cleaves PAR1 at R41 and promotes G protein inflammatory signaling. *B* and *C*, Endothelial EA.hy926 cells were treated with 20 nM APC for various times and Cav1 tyrosine (Y)14 phosphorylation was detected. Total Cav1 was used as a loading control. Data were quantified (mean ± S.D.) from four independent experiments and expressed as the fraction relative to the untreated control (0 min), and analyzed by one-way ANOVA, \**p* < 0.05. *D*, *E* and *F*, cells were pretreated with dasatinib or DMSO prior to addition of APC. Cell lysates were immunoblotted to detect Cav1 Y14 and c-Src Y416 phosphorylation as indicated. *G, H, I* and *J*, Cells transfected with non-specific (NS) or c-Src specific siRNA were treated with or without APC, lysed and immunoblotted as indicated. β-tubulin and GAPDH were used as loading controls. The data were quantified (mean ± S.D.) from four independent experiments and expressed as the fraction relative to the untreated control, and analyzed by Student’s t-test or one-way ANOVA, \**p* < 0.05; \*\**p* < 0.01; \*\*\**p* < 0.001; \*\*\*\**p* < 0.0001.

Src family kinases (SFKs) are known to promote phosphorylation of Cav1 at the Y14 site. To determine if SFKs mediate APC-induced Cav1 Y14 phosphorylation we used dasatinib, a potent phamacological inhibitor of SFKs. In endothelial cells treated with DMSO vehicle, APC induced a marked increase in Cav1 Y14 phosphorylation that was significantly inhibited in cells preincubated with dasatinib (Fig. 1 *D*, top panels, lanes 1-2 vs. 3-4, and *E*). Importantly, APC induced a concomitant increase in Src Y416 phosphorylation (Fig. 1 *D*, lower panels, lanes 1-2, and *F*), which results from Src conformational changes that enables release from an autoinhibited state and facilitates Y416 autophosphorylation (30). APC-stimulated Src Y416 phosphorylation was also blocked by dasatinib in endothelial cells (Fig. 1 *D*, lanes 1-2 vs. 3-4, and *F*). To determine whether c-Src mediates APC-induced Cav1 Y14 phosphorylation, we used an siRNA knockdown approach. Expression of c-Src was virtually abolished in cells transfected with the c-Src siRNA compared to non-specific siRNA control transfected cells (Fig. 1 *G*, lower panels, lanes 1-2 vs. 3-4, and *H*), indicating that siRNA significantly reduced c-Src expression. In contrast to non-specific siRNA transfected cells, APC-induced Cav1 Y14 phosphorylation was significantly inhibited in c-Src depleted cells (Fig. 1 *G*, top panels, lanes 1-2 vs. 3-4, and *I*). As expected, APC-stimulated c-Src Y416 phosphorylation was not detectable in endothelial cells with diminished c-Src expression (Fig. 1 *G*, lower panels, lanes 1-2 vs. 3-4, and *J*). These data suggest that c-Src mediates Cav1 Y14 phosphorylation following APC-activation of PAR1 in endothelial cells.

### βarr2-regulates APC-activated PAR1 induced Cav1 Y14 phosphorylation and c-Src activation

βarr2 functions as a central driver of APC/PAR1 endothelial cytoprotective signaling (3, 10) and β-arrestins have been reported to regulate c-Src activity (31, 32). These studies suggest that βarr2 may be required for c-Src-mediated Cav1 Y14 phosphorylation induced by APC-activation of PAR1. To examine the role of βarr2, endothelial cells were transfected with βarr2-specific siRNA or non-specific siRNA control, treated with APC and Cav1 Y14 phosphorylation determined. In cells deficient in βarr2 expression, basal Cav1 Y14 phosphorylation was increased even in the absence of agonist stimulation compared to non-specific siRNA transfected cells (Fig. 2 *A*, lanes 1 vs. 3 and *B*). Nonetheless, APC failed to increase Cav1 Y14 phosphorylation in βarr2 deficit cells compared to cells transfected with non-specific siRNA (Fig. 2, *A*, lanes 1-2 vs. 3-4, and *B*). These results suggest that βarr2 is important for APC-induced Cav1 Y14 phosphorylation. To determine the function of βarr2 in c-Src activation induced by APC/PAR1 in endothelial cells, c-Src Y416 phosphorylation in βarr2 deficient cells was examined. In non-specific siRNA control cells, APC induced a robust increase in c-Src Y416 phosphorylation that was significantly inhibited in cells lacking βarr2 expression (Fig. 2 *C*, lanes 1-2 vs. 3-4 and *D*). Taken together, these findings suggest that APC-activation of PAR1 promotes Cav1 Y14 phosphorylation via a βarr2- and c-Src-dependent pathway.

**Figure 2.**
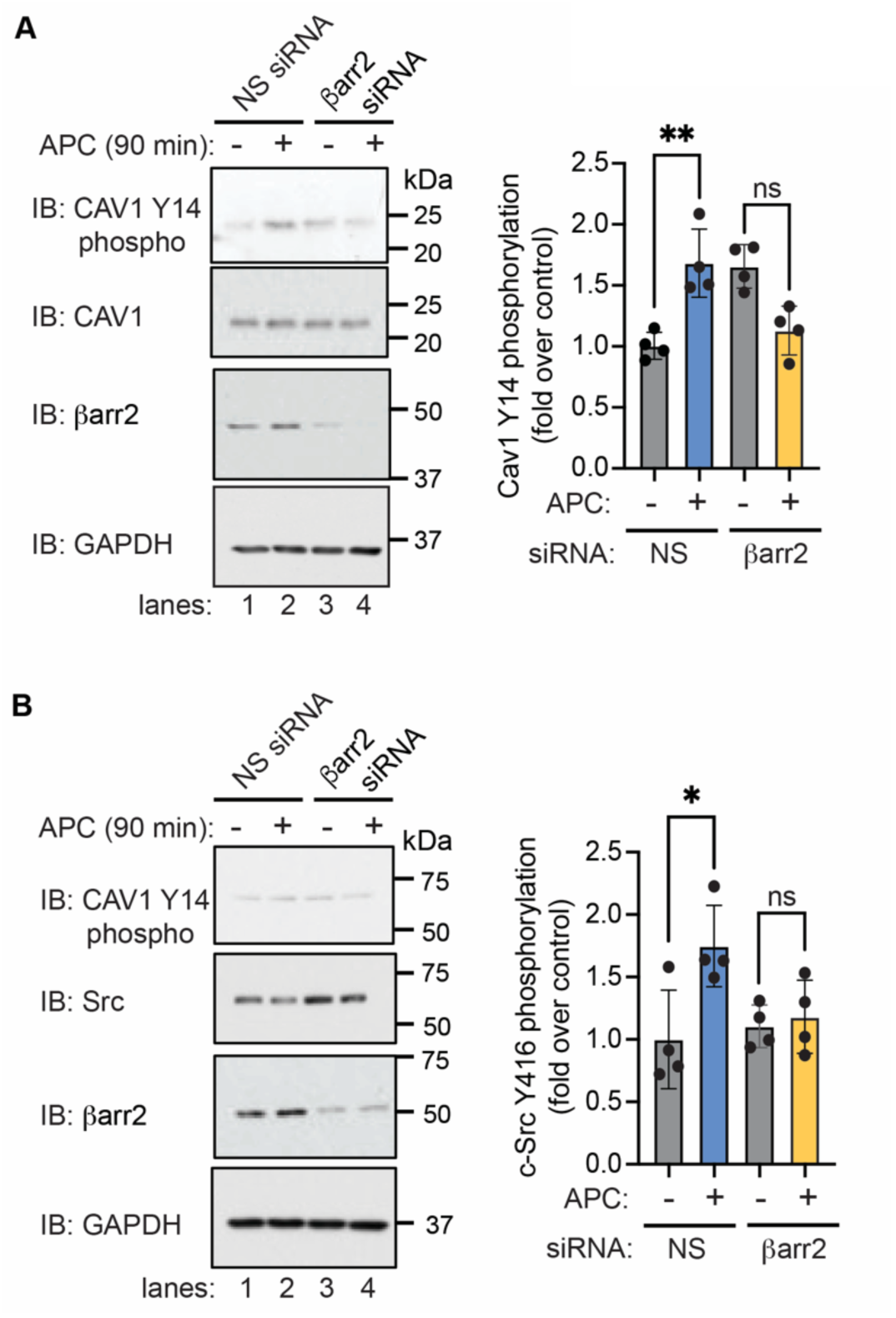
βarr2 is required for APC-stimulated Cav1 Y14 phosphorylatin and c-Src Y416 phosphorylation. Endothelial EA.hy926 cells were transfected with non-specific (NS) or βarr2-specific siRNA, treated with 20 nM APC and Cav1 tyrosine (Y)14 phosphorylation (*A* and *B*), and c-Src Y416 phosphorylation (*C* and *D*), were detected by immunoblotting as indicated. GAPDH was used as a loading control. The data were quantified (mean ± S.D.) from at least four independent experiments and expressed as the fraction relative to the untreated control, and analyzed by one-way ANOVA, \**p* < 0.05; \*\**p* < 0.01.

### APC regulates PAR1-Cav1 association and colocalization revealed by STORM imaging

Next we sought to determine if APC modulates endogenous PAR1 and Cav1 colocalization using STORM combined with total internal reflection fluorescence (TIRF) imaging. STORM enables nanometer-scale (20-30 nm) resolution and was used to visualize endogenous PAR1 and Cav1 single molecules in caveolae at or near the plasma membrane. To validate the specificity of the anti-PAR1 antibody for STORM imaging, we used HEK293 parental cells transiently transfected with human PAR1 or pcDNA3 vector. To validate the anti-Cav1 antibody, we compared HEK293 parental cells with HEK293 CRISPR-Cas9 Cav1/2 knockout cells. In HEK293 cells transiently expressing PAR1, the anti-PAR1 WEDE antibody raised against the N-terminus of PAR1 detected PAR1 at the cell surface and in endocytic vesicles, whereas no signal was observed in pcDNA3-transfected control cells (Fig. 3, *A*). HEK293 parental cells immunostained with anti-Cav1 antibodies recognized discrete caveolae-like puncta enriched in Cav1 predominantly at the cell periphery, which were absent in the Cav1,2 KO cells lacking Cav1 expression (Fig. 3, *B*). The specificity of the anti-Cav1 antibody was further confirmed by immunoblotting lysates prepared from HEK293 parental and Cav1,2 KO cells (Fig. 3, *C*).

**Figure 3.**
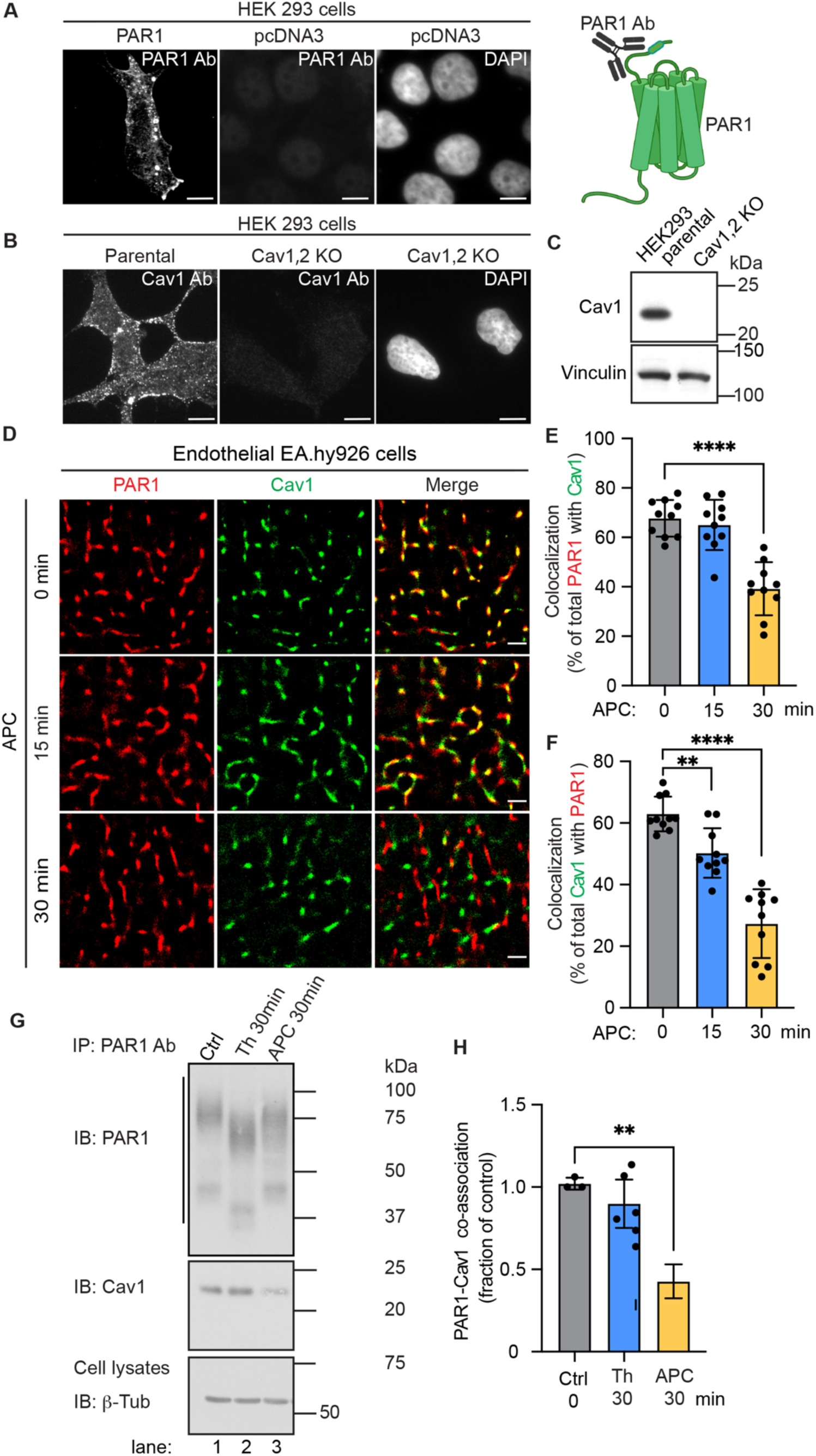
APC modulates endogenous PAR1-Cav1 co-localization assessed by STORM imaging. *A*, HEK293 cells transfected with human PAR1 or pcDNA3 plasmids were immunostained with the anti-PAR1 WEDE antibody (Ab) which recognizes the N-terminus of PAR1 (cartoon). PAR1 expression was detected by immunofluorescence confocal microscopy. Scale bar, 100 μm. *B*, HEK293 parental cells and Cav1,2 knockout (KO) cells were immunostained for Cav1 aind imaged by confocal microscopy. Scale bar, 100 μm. *C*, HEK293 parental cells and Cav1,2 KO cell lysates were immunblotted for Cav1. Vinculin was used as loading control. *D*, Endothelial EA.hy926 cells were incubated with or without APC (20 nM) for various times, processed, immunostained for PAR1 (*red*) and Cav1 (*green*) and imaged by STORM. Scale bar, 1 μm. *E* and *F*, PAR1-Cav1 co-localization was quantified (mean ± S.D.) from three independent experiments using ten cells per condition per experiment, and expressed as a percent of total PAR1 with Cav1 and percent of total Cav1 with PAR1. Data were analyzed by one-way ANOVA, \*\**p* < 0.01; \*\*\*\**p* < 0.0001. *G*, Endothelial EA.hy926 cells treated with or without APC (20 nM) or Th (10 nM) were immunoprecipited (IP) with anti-PAR1 antibodies (Ab) and endogenous PAR1 and Cav1 were detetected by immunoblotting. β-tubulin was used as a loading control. *H*, PAR1-Cav1 co-association was quantified (mean ± S.D.) from three independent experiments and expressed as fraction over untreated control (0 min) and analyzed by ANOVA, \*\**p* < 0.01.

Using these validated antibodies, we then assessed the effect of APC on endogenous PAR1 and Cav1 colocalization in endothelial EA.hy926 cells by STORM imaging. In unstimulated cells, substantial PAR1-Cav1 colocalization was detected, with ∼70% of PAR1 colocalized with Cav1 and ∼60% of Cav1 colocalized with PAR1 (Fig. 3, *D, E* and *F*). APC stimulation for 15 min did not alter the extent of PAR1–Cav1 colocalization, which remained comparable to untreated controls (Fig. 3, *D, E* and *F*). However, a significant decrease in PAR1-Cav1 colocalization was observed following APC stimulation at 30 min and was reflected by a reduction from ∼70% to ∼40% for PAR1-Cav1 colocalization and ∼60% to ∼30% for Cav1-PAR1 colocalization (Fig. 3, *D, E* and *F*). These results are consistent with APC-induced disruption of PAR1-Cav1 protein complexes.

To substantiate PAR1-Cav1 protein-protein interaction in endothelial cells we used co-immunoprecipitation and examined whether PAR1-Cav1 co-association is regulated by APC or thrombin. Endothelial cells were either left untreated or treated with thrombin or APC, PAR1 was immunoprecipitated with anti-PAR1 antibodies and co-associated Cav1 was detected. In control cells, a substantial fraction of PAR1 co-associated basally with Cav1 (Fig. 3, *G* lane 1, and *H*). After thrombin incubation, cleaved and activated PAR1 migrated as lower molecular weight species and retained binding to Cav1 comparable to that observed in untreated control cells (Fig. 3, *G* lanes 1-2 and *H*). In contrast to thrombin, APC-activated PAR1 exhibited a more modest shift in size of the cleaved receptor but showed a significant disruption of the PAR1-Cav1 complex resulting in a significant loss of Cav1 association with PAR1 (Fig. 3, *G* lanes 2 vs. 3 and *H*). Thus, these data collectively, suggest that APC activation of PAR1 induces Cav1 Y14 phosphorylation and disrupts Cav1 association with activated PAR1.

### Endogenous GRK5 localization to caveolae is modulated by APC stimulation

GRK5 regulates APC-activated PAR1 induced βarr2 recruitment (28) and GRK5 has previously been shown to directly interact with Cav1 *in vitro* using purified proteins (27). However, it is not known if endogenous GRK5 resides in caveolae in intact cells and was examined. We first used sucrose density gradient fractionation where detergent resistant membranes enriched in Cav1 separate to lower density fractions. GRK5 was detected in both the Cav1-enriched fractions and heavier fractions containing the early endosome antigen-1 (EEA1) peripheral membrane protein (Fig. 4, *A*). Thus, like PAR1 (3, 10), GRK5 is localized both within and outside of Cav1-enriched membrane microdomains in endothelial cells.

**Figure 4.**
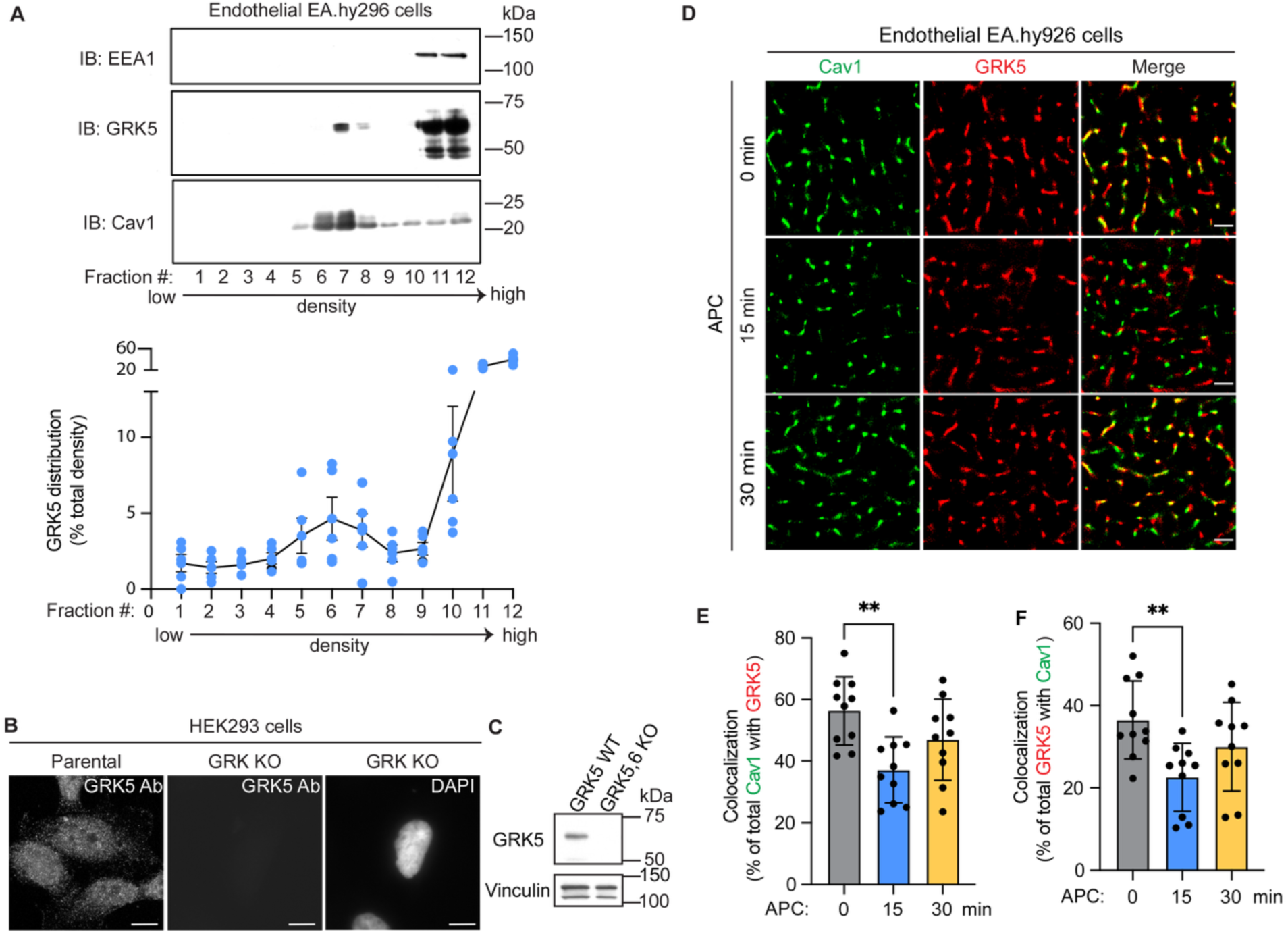
Endogenous GRK5 localizes to plasma membrane caveolae microdomains in endothelial cells. *A*, Endothelial EA.hy926 cell lysates were subjected to sucrose density gradient fractionation and collected fractions were immunoblotted for Cav1, GRK5 and early endosomal antigen-1 (EEA1) as indicated. The data (mean ± S.D.) from six independent experiments are expressed as the percentage GRK5 in each fraction compared to the total combined GRK5 signal intensity across all fractions. *B*, specificity of the GRK5 antibody was validated by immunofluorescence using HEK293 parental and CRISPR-Cas9 GRK KO cells. Scale bar, 100 μm. *C*, Immunoblot of lysates from HEK293 parental and GRK5,6 KO cells confirming GRK5 antibody specificity. Vinculin was used as a loading control. *D*, EA.hy926 cells were treated with APC for various times, processed, immunostained for Cav1 (*green*) and GRK5 (*red*) and imaged by STORM. Scale bar, 1 μm. *E* and *F*, Cav1-GRK5 co-localization was quantified (mean ± S.D.) from three independent experiments using ten cells per condition per experiment, and expressed as a percent of total Cav1 colocalized with GRK5 and percent of total GRK5 colocalized with Cav1. Data were analyzed by one-way ANOVA, \*\**p* < 0.01.

Next, we used STORM imaging to determine if APC regulates endogenous GRK5 localization in caveolae in endothelial cells by direct visualization. The anti-GRK5 antibody was validated in HEK293 CRISPR-Cas9 GRK KO cells lacking GRK2,3 and 5,6 isoforms using immunofluorescence confocal microscopy. The antibody detected GRK5 in parental cells but GRK5 detection was absent in GRK KO cells (Fig. 4, *B*). To further assess antibody specificity, we used lysates from HEK293 CRISPR-Cas9 GRK5,6 KO cells and compared GRK5 expression to HEK293 parental cells by immunoblot analysis. Consistent with confocal imaging, the antibody detected GRK5 expression in HEK293 parental cells but not in the GRK5,6 isoform specific KO cell lysates (Fig. 4, *C*), suggesting that the antibody specifically and reliably detects the GRK5 protein, as previously reported (33). In the absence of APC stimulation, STORM imaging detected endogenous Cav1 and GRK5 single molecules in endothelial cells with substantial colocalization at 0 min with ∼58% of Cav1 colocalized with GRK5 and ∼38% of GRK5 colocalized with Cav1 (Fig. 4, *D, E* and *F*). After stimulation with APC for 15 min, a significant decrease in total Cav1-GRK5 and GRK5-Cav1 colocalization was observed and indicated by a reduction from 58% to 38% for Cav1-GRK5 colocalization and 38% to 22% for GRK5-Cav1 colocalization (Fig. 4, *D*, *E* and *F*). However, after 30 min of APC stimulation the extent of Cav1-GRK5 and Cav1-GRK5 colocalization returned to levels comparable with that observed basally in unstimulated cells (Fig. 4, *D*, *E* and *F*). These findings suggest that APC dynamically regulates GRK5-Cav1 association in caveolae plasma membrane microdomains, prior to disruption of PAR1-Cav1 protein-protein interaction.

### The GRK5 Cav1 binding motif regulates localization to the plasma membrane

GRK5 binds directly to Cav1 *in vitro* through an N-terminal region containing a consensus Cav1 binding aromatic-rich motif <u>I</u>^62^GRLL<u>F</u>RQ<u>F</u>^70^ that binds to the caveolin-1 CSD (27) (Fig. 5, *A*). To determine whether the Cav1 binding motif of GRK5 is important for GRK5-Cav1 interaction in intact cells, a GRK5 mutant containing conserved aromatic and hydrophobic isoleucine (I)62, phenylalanine (F)67 and F70 residues were converted to alanine (A) was generated and termed the GRK5 “IFF” mutant. We first examined GRK5 IFF mutant expression and localization by immunofluorescence confocal microscopy in HeLa cells. GRK5 is known to localize to the plasma membrane via a C-terminal amphipathic helix and was found predominantly at the plasma membrane in HeLa cells (Fig. 5, *B*). Unexpectedly we found that the GRK5 IFF mutant localized primarily in the cytoplasm similar to a previously reported GRK5 amphipathic helix 4A mutant harboring leucine (L) to alanine (A) mutations at positions L550A, L551A, L554A, F555A (24) (Fig. 5, *B*). To verify GRK5 IFF mis-localization from the plasma membrane we utilized a high-speed ultracentrifugation cell fractionation assay capable of separating soluble/cytosolic proteins from particulate/membrane bound proteins. A significantly greater fraction of GRK5 WT expressed in HEK293 cells was detected in the particulate membrane fraction compared to the soluble fraction, similar to the modestly expressed endogenous GRK5 detected in these cells (Fig. 5 *C*, lanes 1-2 vs. 3-4.). However, both the GRK5 IFF and 4A mutant showed a greater distribution to the soluble cytosolic fraction compared to GRK5 WT (Fig. 5 *C*, lanes 5-8 vs. 3-4.). These results suggest that the GRK5 IFF mutant is mis-localized and resides predominantly in the cytoplasm like the previously reported GRK5 4A mutant (24).

**Figure 5.**
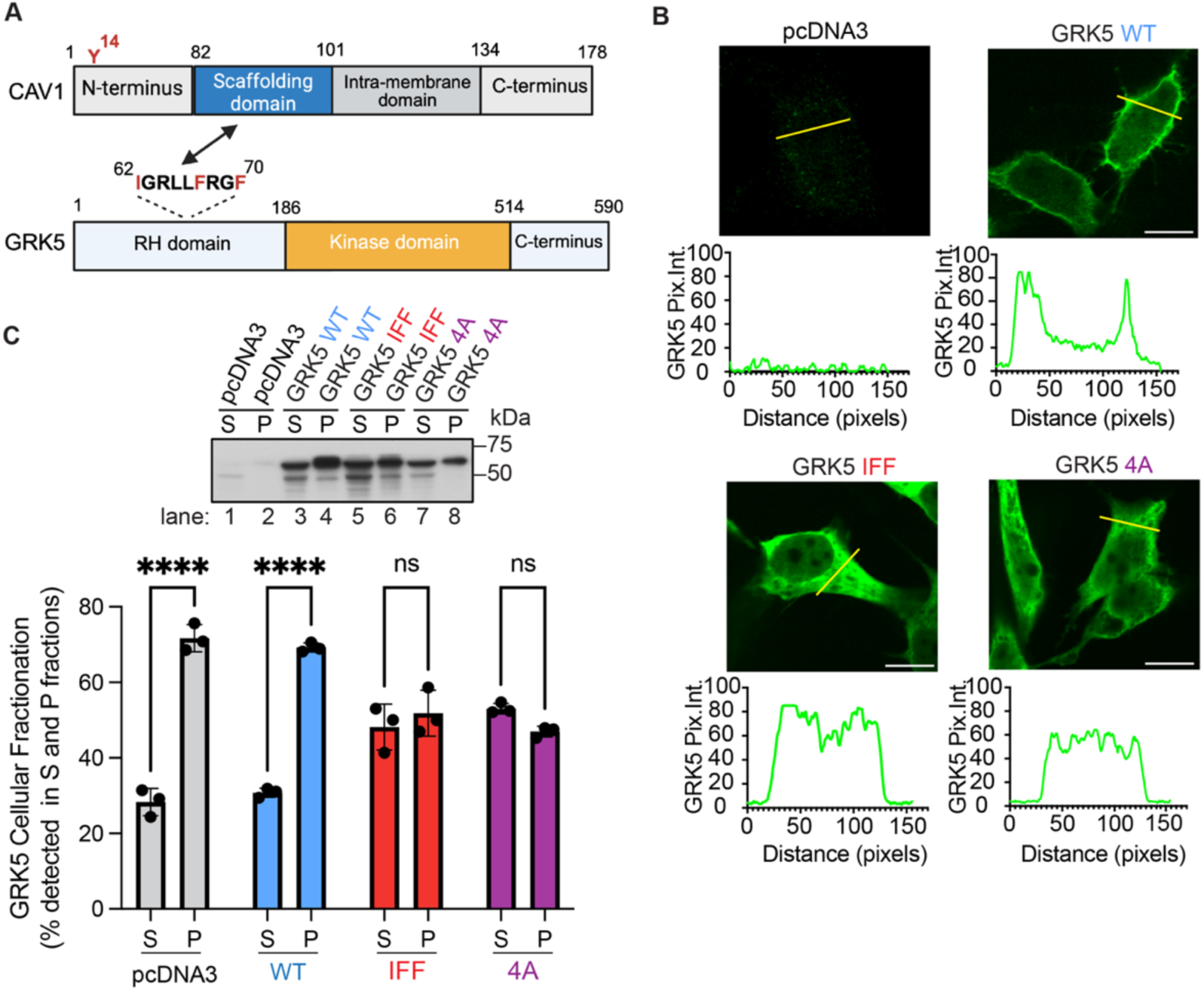
The GRK5 Cav1 binding IFF motif is important for membrane localization. *A*, Schematic of Cav1 (amino acids 1-178) and GRK5 (amino acids 1-590) protein domains. Cav1 contains a N-terminus tyrosine (Y)14 residue, Cav1 scaffolding domain (CSD), intra-membrane domain and C-terminus. GRK5 contains a consenses Cav1 binding motif (residues 62–70) within the RH domain, and a kinase domain and C-terminus. The putative Cav1 binding motif interaction with Cav1 scaffolding domain is indicated by the double-headed arrow. The GRK5 IFF residues mutated to alanine are highlighted in red. *B*, HeLa cells transiently expressing GRK5 WT, IFF or 4A mutants were immunostained and imaged by confocal microscopy. Scale bar, 100 μm. GRK5 subcellular localization was quantified by line scan analysis of pixel intensity using ImageJ software. *C*, GRK5 WT, IFF, 4A mutant or pcDNA were transfected into HEK293 cells, and subjected to cellular fractionation resulting in a soluble (S)/cytosolic fraction and a particulate (P)/membrane fractions. GRK5 was detected by immunoblotting and quantified. The percent of GRK5 detected in S and P fractions was determined from three independent experiments and analyzd by one-way ANOVA, \*\*\*\**p* < 0.0001.

### A consensus Cav1 binding motif is required for GRK5-Cav1 interaction in cells

Next, we examined the interaction of GRK5 wild-type and IFF mutant with Cav1 in HEK293 cells by co-immunoprecipitation. Wildtype GRK5 robustly co-associated with HA-tagged Cav1 in immunoprecipitates compared to the IgG control IPs (Fig. 6, *A*, lanes 1 and 2 and *B*). In contrast to GRK5 WT, the GRK5 IFF mutant showed a significant reduction in Cav1 interaction in immunoprecipitates (Fig. 6, *A*, lanes 2 and 3, and *B*), suggesting that the conserved Cav1 binding IFF motif is critical for GRK5–Cav1 interaction in intact cells. To determine whether the loss of GRK5 membrane anchoring observed with the GRK5 IFF mutant contributed to the loss of Cav1 binding, we examined Cav1 interaction with the GRK5 4A mutant. In contrast to the GRK5 IFF mutant, GRK5 4A retained the capacity to interact with HA-Cav1 like GRK5 WT despite altered localization primarily to the cytoplasm (Fig. 6, *C*, lanes 3-5 vs. 1-2 and *D*). Thus, the loss of GRK5 membrane anchoring is not sufficient to disrupt GRK5-Cav1 interaction in cells.

**Figure 6.**
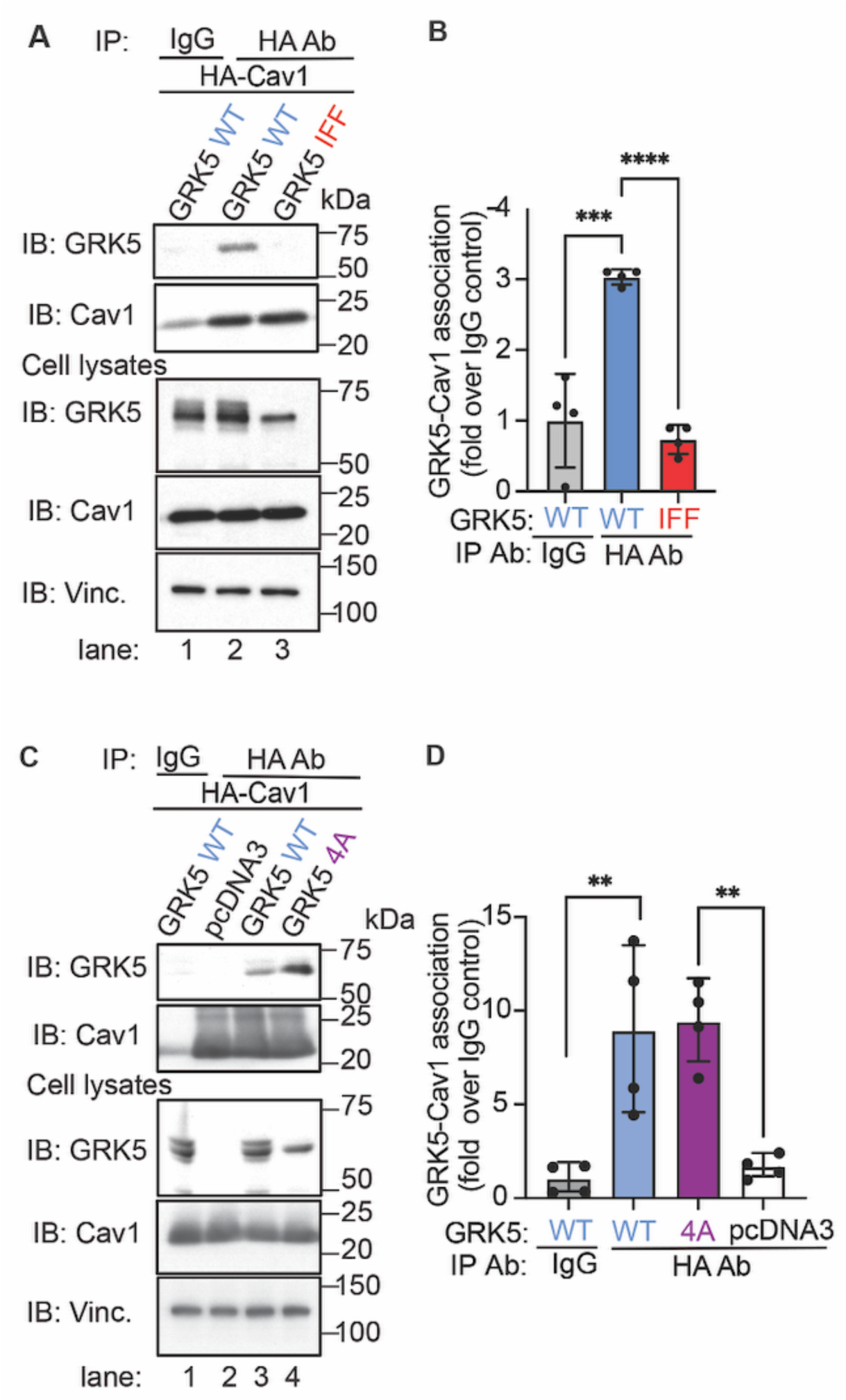
The GRK5 IFF motif mediates interaction with Cav1 in intact cells. *A* and *C*, HEK293 cells transiently expressing HA-Cav1 together with either GRK5 WT, mutants or pcDNA3 were processed, immunoprecipitated and immunoblotted as indicated. Cell lysates were immunoblotted for total HA-Cav1, GRK5 and Vinculin expression. *B* and *D,* the data (mean ± S.D.) from four independent experiments are expressed as fold over IgG control and were analyzed by one-way ANOVA, \*\**p* < 0.01; \*\*\**p* < 0.001; \*\*\*\**p* < 0.0001.

### GRK5 wildtype and neither the IFF nor 4A mutant support APC-induced βarr2 recruitment to activated PAR1

APC-activated PAR1 requires GRK5 for recruitment of βarr2 (28) and βarr2 drives endothelial cytoprotection (3, 10). However, it is not known whether GRK5-Cav1 interaction is important for APC/PAR1-induced βarr2 recruitment and was examined using bioluminescence resonance energy transfer (BRET) assays. In these studies, HEK293 cells co-expressing PAR1 fused to yellow fluorescent protein (YFP), the APC co-receptor EPCR and βarr2-fused the *Renilla* luciferase (Rluc) were stimulated with APC and βarr2 recruitment was monitored by changes in BRET (Fig. 7, *A*). APC induced a robust hyperbolic increase in βarr2 recruitment measured as an increase net BRET in HEK293 parental cells (Fig. 7, *A*), whereas the APC proteolytically inactive variant containing serine (S)360 to alanine mutation failed to promote βarr2 recruitment (Fig. 7, *B*). Thus, the HEK293 model system recapitulates the requirement for APC cleavage of PAR1 to induce βarr2 recruitment. (34).

**Figure 7.**
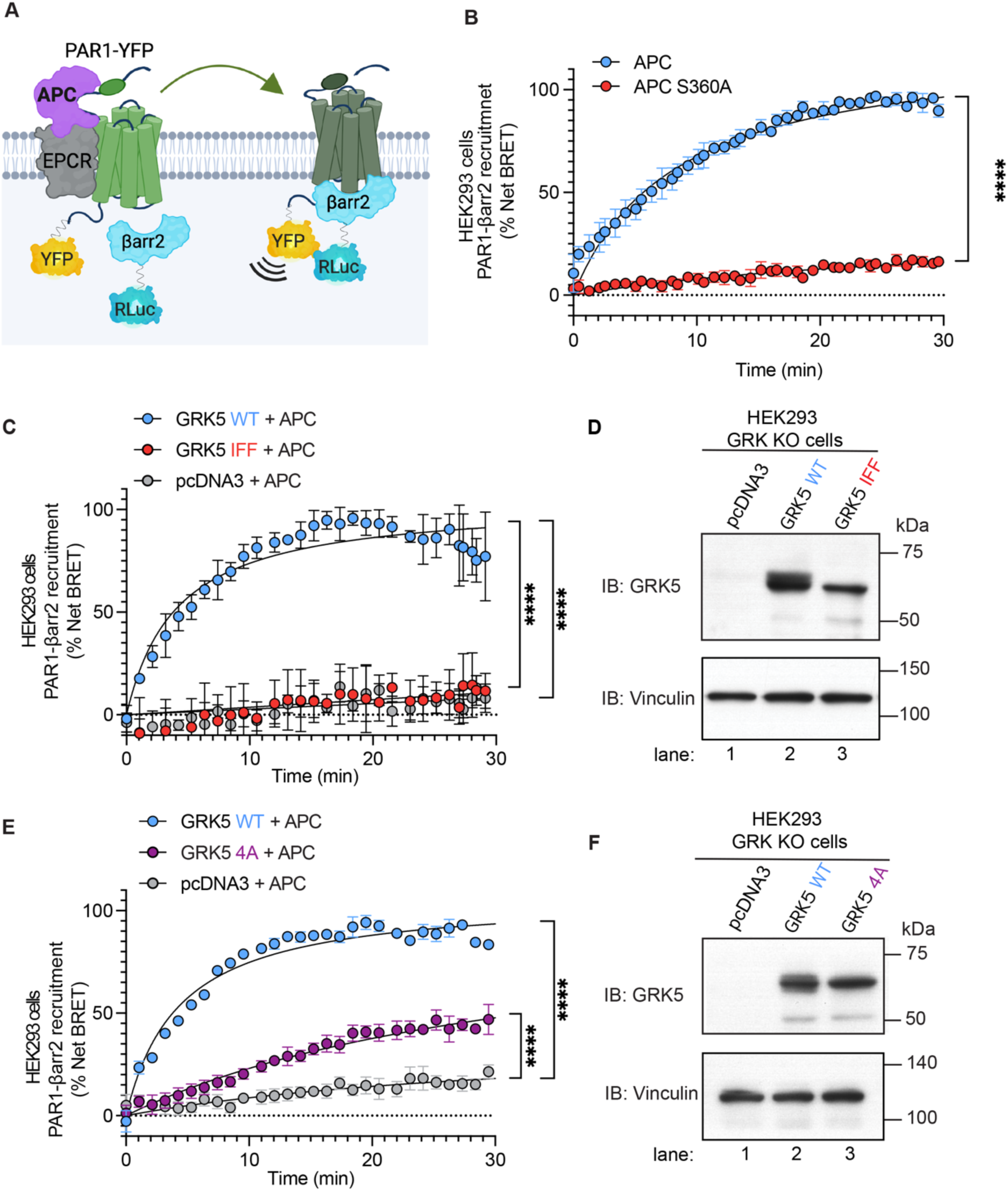
GRK5 wildtype but niether the GRK5 IFF nor 4A mutant support APC-activated PAR1 induced βarr2 recruitment. *A*, Cartoon, APC bound to EPCR activates PAR1-YFP resulting in the recruitment of Rluc-βarr2 in the BRET assay. *B*, HEK293 cells transiently expression PAR1-YFP, EPCR-Halo and Rluc-βarr2 were treated with 20 nM APC or APC S360A proteolytically inactive mutant and net BRET determined. Data (mean ± S.D.) from three independent experiments were analyzed by one-way ANOVA, \*\*\*\**P* < 0.0001. HEK293 CRISPR-Cas9 GRK knockout (KO) cells transiently expressing PAR1-YFP, EPCR-Halo, Rluc-βarr2 and either GRK5 wildtype (WT), GRK5 IFF mutant or pcDNA3 (*C* and *D*) or GRK5 WT, GRK5 4A mutant or pcDNA3 (*E* and *F*) were stimulated with 20 nM APC and net BRET determined. Cell lysates were immunoblotted for GRK5 and vinculin expression. Data (mean ± S.D.) from three or four independent experiments were analyzed by one-way ANOVA, \*\*\*\**p* < 0.0001.

Next, we used HEK293 CRISPR-Cas9 GRK KO cells expressing PAR1-YFP, the APC co-receptor EPCR and βarr2-Rluc together with GRK5 wildtype, IFF and 4A mutant and assessed the effect on APC-induced βarr2 recruitment using BRET. In cells co-expressing GRK5 wildtype, APC stimulated a robust increase in recruitment of βarr2 to activated PAR1 (Fig. 7, *C* and *D*, lane 2), whereas APC failed to stimulate βarr2 recruitment in pcDNA3 vector transfected cells (Fig. 7, *C* and *D*, lane 1). These findings indicate that GRK5 expression is critical for APC-induced βarr2 recruitment. In contrast to wildtype GRK5, expression of the GRK5 IFF mutant failed to restore APC-induced βarr2 recruitment to activated PAR1 (Fig. 7, *C* and *D*, lane 3). Similarly, in cells expressing the GRK5 4A mutant, APC-stimulated a modest quasi-linear increase in βarr2 recruitment (Fig. 7, *E* and *F*, lanes 1-3), consistent with a critical function for GRK5 membrane anchoring in facilitating APC-induced βarr2 recruitment as recently reported (28). These results indicate that although GRK5-Cav1 interaction may contribute to βarr2 recruitment, proper GRK5 membrane localization is likely most critical for APC/PAR1-induced βarr2 recruitment.

## Discussion

In this study we show that APC regulates Cav1 function by modulating Cav1 Y14 phosphorylation and Cav1 association with both PAR1 and GRK5 that likely contribute to APC-activated PAR1 βarr2 recruitment. We previously demonstrated that Cav1 expression is critical for APC/ PAR1-induced endothelial cytoprotection (10, 16). However, the mechanisms underlying this requirement is not known. Here we report that APC-activation of PAR1 promotes Cav1 Y14 phosphorylation via a βarr2- and c-Src-dependent pthaway. In addition, we report that endogenous PAR1-Cav1 basally associate and that APC activation of PAR1 disrupts PAR1-Cav1 complexes in endothelial cells. We also found that endogenous GRK5 localizes to caveolae in endothelial cells and GRK5-Cav1 association is dynamically regulated by APC stimulation of PAR1. Our results further demonstrate that mutation of the GRK5 caveolin-1 binding motif disrupts interaction with Cav1 in intact cells, perturbs membrane association and fails to support APC-activated PAR1 induced βarr2 recruitment, in contrast to wildtype GRK5.

Caveolae are cholesterol rich, highly abundant specialized lipid rafts in endothelial cells and are essential for APC-activated PAR1 cytoprotective signaling. The formation of caveolae in endothelial cells requires Cav1, a principal structural component, and cholesterol (17). Several studies showed that depletion of cholesterol using methyl-β-cyclodextrin, which disrupts both lipid rafts and caveolae impairs APC/PAR1 signaling in endothelial cells (15, 16). To assess the role of caveolae more directly, a Cav1 shRNA or siRNA targeting strategy was taken and shown to abolish Cav1 expression without perturbing surface expression of PAR1 or EPCR, the APC cofactor, and blocked APC/PAR1-mediated endothelial barrier protection and anti-apoptotic responses (10, 16). Additionally, sucrose density gradient fractionation was used to assess PAR1 localization in Cav1 enriched membranes (10, 16), indicating that PAR1 and Cav1 are present in the same dense fractions. However, direct visual evidence that endogenous PAR1 localizes to caveolae in intact endothelial cells had not been directly demonstrated. Caveolae are small plasma membrane invaginations (50-100 nm) and requires super resolution imaging for direct visualization (17). Here we used STORM imaging to visualize caveolae and demonstrate that a substantial population of PAR1 molecules colocalize with Cav1 molecules, providing strong evidence that endogenous PAR1 resides in caveolae in endothelial cells.

Cav1 Y14 phosphorylation induces conformational changes that spatially separate Cav1 molecules and may alter the conformation or accessibility of the CSD (29, 35). The Cav1 CSD mediates protein-protein interactions through recognition of a caveolin binding motif originally identified as aromatic-rich sequences that contain a specific spacing (ØXØXXXXØ, ØXXXXØXXØ, or ØXØXXXXØXXØ, where Ø is Trp, Phe, or Tyr), as originally reported (36). The CSD protein-protein interactions are also regulated by Cav1 Y14 phosphorylation, which has been shown to enhance Cav1 CSD-focal adhesion interactions (37) and increase Cav1 CSD-endothelial nitric oxide synthase (eNOS) association (38, 39). In a rodent animal model, phosphorylation of Cav1 was also shown to enhance GRK2-Cav1 interaction and modulate endothelial nitric oxide synthase expressed in sinusoidal endothelial cells (40). We found that APC induces Cav1 Y14 phosphorylation through a βarr2 and c-Src dependent pathway and promotes disruption of PAR1-Cav1 complexes. However, PAR1 appears to lack a caveolin binding motif in its’ cytoplasmic regions and not likely to directly interact with Cav1. We therefore propose that an intermediary protein with a consensus caveolin binding motif most likely regulates PAR1-Cav1 association following APC stimulation that remains to be identified.

A critical role for GRK5 in APC/PAR1-induced βarr2 recruitment has been recently reported (28). Interestingly, the N-terminus of GRK5 contains a consensus caveolin binding motif that directly interacts with the Cav1 CSD *in vitro* and potently inhibits GRK5 activity (27). Whether endogenous GRK5 is present in caveolae and interacts with Cav1 in intact cells is not known and was examined. In this study we found that endogenous GRK5 is present in caveolin-1 enriched fractions using sucrose density gradient fractionation and directly visualized to colocalize with caveolin-1 basally by super resolution STORM imaging. Thus, endogenous GRK5 localizes to caveolae in endothelial cells. In addition, APC-stimulation of PAR1 dynamically regulates GRK5-Cav1 interaction causing GRK5-Cav1 dissociation and re-association within 30 min. We hypothesize that disruption of the Cav1-GRK5 complex may permit APC-induced GRK5 activation and phosphorylation of PAR1, which is critical for βarr2 recruitment as recently report (28).

GRK5 is primarily anchored at the plasma membrane via a C-terminus amphipathic helix (23–25). Unexpectedly, we found that the GRK5 IFF mutant, which lacks interaction with caveolin-1 is also defective in membrane association, whereas the GRK5 4A amphipathic helix mutant localizes primarily in the cytoplasm but retained Cav1 binding. These findings indicate that additional determinants of plasma membrane localization are perturbed in the GRK5 IFF mutant. In addition to the C-terminus amphipathic helix, the GRK5 N-terminal basic region binds to the plasma membrane lipid phosphatidylinositol 4,5-bisphosphate (41, 42)and may contribute to plasma membrane localization. Moreover, GRK5 dimerization mediated by an N-terminal short sequence may form a hydrophobic dimeric interface that may facilitate multivalent interaction with plasma membrane lipids enhancing plasma membrane localization. Whether the GRK5 IFF mutant affects phosphatidylinositol 4,5-bisphosphate lipid binding or dimerization is not known.

In this study we show that APC modulates Cav1 Y14 phosphorylation through a βarr2-dependent c-Src pathway and dynamically regulates PAR1-Cav1 and GRK5-Cav1 coassociation in caveolae. We further show that GRK5-Cav1 interaction occurs through an N-terminal Cav1 binding motif in intact cells and a mutant GRK5 defective in Cav1 binding and plasma membrane localization fails to support APC-activated PAR1-induced βarr2 recruitment. Moreover, APC proteolytic activity is required to induce βarr2 recruitment to activated PAR1, consistent with our previous findings showing that an active site–blocked APC-dansyl-EGR chloromethyl ketone signaling in endothelial cells is markedly inhibited compared to full active APC (16). Thus, APC cleavage and activation of PAR1 βarr2 biased signaling is regulated by Cav1 through multiple mechanisms that likely converve on GRK5. Our findings identify a novel role of Cav1–GRK5 interactions in regulating APC/PAR1 biased βarr2 signaling. Given the importance of βarr2-mediated cytoprotection in endothelial cells, this pathway may represent a therapeutic target for vascular disorders such as sepsis or thrombosis.

## Experimental Procedures

### Cell culture

EA.hy926 cells (ATCC, #CRL-2922) were cultured as previous described (10). Briefly, cells were grown at 37°C, 8% CO_2_ in Dulbecco’s Modified Eagle Medium (DMEM) from Gibco (#10–013-CV and #10437–028) supplemented with 10% fetal bovine serum (FBS) and 20% preconditioned media. HEK293T cells, HEK293A parental, GRK KO, GRK5,6 KO cells (from Dr. Asuka Inoue, Tohoku University) and HEK293 Cav1,2 KO cells (from Dr. Tracy Handel, UC San Diego) were cultured in DMEM supplemented with 10% fetal bovine serum (FBS) (vol/vol) with 5% CO2 atmosphere at 37°C. HeLa cells were cultured in DMEM supplemented with 10% fetal bovine serum (FBS) and 250 μg/mL hygromycin for maintenance as previously described (43).

### Antibodies

In this study, the following antibodies were used: mouse anti–PAR1 WEDE (Beckman Coulter, #IM2584), anti-Cav1 (CST, #3267S and BD, #610060), anti-Cav1 Y14 phospho antibody (CST, #3251), anti-βarr2 (Abcam, #ab54790), GAPDH (GeneTex, #GT239), c-Src Y416 (CST, #2101), anti-c-Src (CST, #2109), GRK5 (Santa Cruz, #SC518005), anti-GRK4-6 (Millipore, #05-466), anti-HA (CST, #3724S), anti-rabbit IgG (CST, #2729), anti-β-Tubulin (CST, #86298), anti-EEA1 (BD Biosciences, #610457), anti-Vinculin (Sigma, #V9131) and anti-HA conjugated to HRP (Roche, #11667475001). The following secondary antibodies were used: anti-mouse or anti-rabbit HRP–conjugated antibodies (Bio-Rad, #170–6516 and #170–6515, respectively), anti-mouse Alexa-488 Fluor (Invitrogen, #A-11001), anti-rabbit Alexa-488 Fluor (Invitrogen, #A-11034), anti-rabbit Alexa-594 Fluor (Invitrogen, #A-11012), anti-mouse Alexa-594 Fluor (Invitrogen, #A-11032), anti-rabbit Alexa-647 Fluor (Invitrogen, #A-21244), and anti-mouse Alexa-647 Flour (Invitrogen, # A-21235) antibodies.

### Agonists and inhibitors

The agonists used in these studies were human APC (Prolytic Haematologic Technologies LLC, #HCAPC-0080), human APC S360A proteolytically inactive mutant was kindly provided by Professor John Griffin (The Scripps Research Institute, La Jolla, CA), human α-thrombin (Enzyme Research Laboratories, #HT 1002a), and dasatinib was purchased from Sigma (#SML2589). Cells were treated with 50 nM dasatinib for 30 min prior to addition of APC.

### Plasmids

Human FLAG-tagged PAR1 cDNA was previously described (44). GRK5 wildtype and 4A mutant in pcDNA3 were generously provided by Dr. Philip Wedegaertner (Thomas Jeffereson University, Philadelphia, PA). The GRK5 IFF mutant, PAR1-YFP, EPCR-Halo, RLuc-βarr2 were generated by Gibson assembly homologous recombination (Gibson Assembly® Master Mix, New England Biolabs) followed by whole plasmid sequencing. Cav1 containing a carboxyl terminal HA-epitope tag (HA-Cav1) was a gift from Ari Helenius (Addgene plasmid #27703; http://n2t.net/addgene:27703; RRID:Addgene_27703) (45).

### SiRNA tranfections

The following siRNAs were used in the study including c-Src siRNA Hs_SRC_11 5’ - GGCGCGGCAAGGTGCCAAATT-3’, custom βarr2 siRNA #666 5’ - GGACCGCAAAGTGTTTGTG-3’ and AllStars negative control non-specific siRNA 5′-CUACGUCCAGGAGCGCACC-3′ and purchased from Qiagen. Endothelial EA.hy926 cells were seeded at 1.5 × 10^5^ cells per well in a 12-well plate, grown overnight, and transfected the next day with siRNAs using the TransIT-X2. After transfections, cells were starved overnight in DMEM containing 0.4% FBS, cells were then washed and starved for 1 h with DMEM supplemented with BSA 1 mg/mL, 10 mM HEPES and 2.8 mM CaCl_2_. Cells were then left untreated or treated with 20 nM APC for 90 min, cell lysates were collected in 2×Laemmli sample buffer (LSB) supplemented with 200 mM of dithiothreitol (DTT). Equivalent amounts of cell lysates were the immunoblotted as indicated and quantified by densitometry using ImageJ software (NIH).

### Immunoprecipitation and immunoblotting

Immunoprecipitation (IP) was carried out as described (10). Briefly, endothelial EA.hy926 cells were seeded at 4.95 × 10^6^ cells per 10-cm dish, grown overnight and then serum starved for 1 h prior to treatment with 20 nM APC or 10 nM thrombin. Cells were lysed with Triton lysis buffer (50 mM Tris pH 7.4, 100 mM NaCl, 5 mM EDTA, 1% v/v TritonX-100, 50 mM NaF, and 10 mm NaPP) supplemented with protease inhibitors, clarified by centrifugation, and protein concentrations determined by bicinchoninic acid (BCA) (Thermo Fisher Scientific, #23221). Equivalent amounts of lysates were then precleard with Protein A-Sepharose (Sigma, #GE17-0780-01) and then incubated with Protein A beads and either IgG or anti-PAR1 WEDE antibody incubated at 4°C overnight. IPs were collected, washed and elucted in 2 x Laemmli Sample Buffer (LSB) containing 200 mM DTT and resolved by SDS-PAGE, transferred to PVDF membranes, immunoblotted and developed by chemiluminescence and quantified by densitometry using ImageJ software. Quantification of changes in protein phosphorylation were normalized to the corresponding total protein of the same sample and expressed as a fraction of the indicated control group. Changes in specific protein expression was determined by normalizing to a specific loading control protein and expressed as a fraction relative to the indicated control group.

HEK293 cells were seeded at 1 x 10^6^ per 6-cm dishes and grown overnight, and transfected with either 1 μg of either GRK5 WT, IFF or 4A mutant or pcDNA3 cDNA plasmids along with 1 μg HA-tagged Cav1 using polyethylenimine (PEI). Cell lysates were clarified by centrifugation for 20 min, quantified by BCA and equivalent amounts of lysates were precleared and incubated with Protein A beads together with IgG or anti-HA antibody at 4°C overnight. IPs were collected, washed and eluted in 2xLSB containing 200 mM DTT, resolved by SDS-PAGE, transferred to PVDF membranes, immunoblotted and visualized as indicated above and quantified by densitometry using ImageJ software.

### Immunofluorescence confocal microscopy

HeLa cells were seeded on 12-mm coverslips precoated with 1-5 μg/cm^2^ fibronectin (Sigma, F1141) in 24-well plates at a density of 0.2 ×10^6^ cells per well and grown overnight at 37°C. Cells were then transfected with either 1 µg of GRK5 WT, IFF or 4A mutants or pcDNA3.1 cDNA plasmids. After 24 h, cells were washed with ice-cold PBS, fixed with 4% PFA for 5 min, permeabilized with cold MeOH for 30 sec and immunostained with anti-GRK5 antibody (Santa Cruz, #SC518005) at 1:100 and followed by anti-mouse Alexa-488 antibody. Coverslips were mounted with ProLong Gold Antifade Mountant (Invitrogen, #P10144). Confocal images of 0.20-μm x–y sections were acquired sequentially using an Olympus IX81 DSU spinning-disk microscope (Tokyo, Japan) equipped with a 60x Plan Apo objective lens (1.4 NA), along with appropriate excitation-emission filters and a Cool SNAP HQ2 CCD camera (Andor) using Metamorph version 7.7.4.0 software (Molecular Devices). Image line scan analysis was performed using Fiji ImageJ.

To validate antibodies for STORM imaging HEK293 cells were seeded on coverslips in 24-well plates at a density of 0.1×10^6^ cells per well and transfected with 1 µg of PAR1 in pcDNA3 or pcDNA3 only. Cells were fixed as described above and then immuostained with anti-PAR1 WEDE antibody followed by secondary anti-mouse Alexa-488 antibody. The Cav1 and GRK5 antibodies were validated using HEK293 parental cells and CRISPR-Cas9 GRK KO, GRK5,6 KO or Cav1,2 KO cells grown on coverslips in 24-well plates at a density of 0.2×10^6^ cells per well using the Cav1 antibody (CST, #3267S) or GRK5 antibody (Santa Cruz, #SC518005) followed by secondary anti-rabbit Alexa-488 antibody or anti-mouse Alexa-488 antibodies, mounted and imaged by confocal microscopy as described above.

### STORM imaging and analysis

STORM imaging was used to analyze co-localization between endogenous PAR1-Cav1 and GRK5-Cav1 molecules in endothelial cells using the validated antibodies. Cells plated in 35-mm glass-bottom dishes were grown in DMEM containing 0.4% FBS overnight and serum starved in DMEM containing 1% BSA and 20 mM HEPES for 1 h at 37°C. Cells were then treated with or without 20 nM APC as indicated. Cells were fixed with 4% PFA, permeabilized and incubated with either anti-PAR1 or anti-GRK5 antibodies together with anti-Cav1. Cells were then incubated with either mouse or rabbit Alexa Flour-488 antibodies followed by anti-Alex Fluor 647-secondary antibodies to detect Cav1 respectively. Cells were the immersed in STORM imaging buffer (50 mM Tris, pH 8.0, 10 mM NaCl, 10% glucose, 0.1M cysteamine (#30070-50G, Sigma-Aldrich),

0.56 mg/mL glucose oxidase (#G2133-250KU, Sigma-Aldrich), and 0.68 mg/mL catalase (#C40-100MG, Sigma-Aldrich). STORM imaging was conducted using a Nikon A1R Confocal STORM microscope system equipped with a 100×/1.49 NA TIRF oil-immersion objective and an ANDOR iXon3 Ultra DU897 EMCCD camera (UC San Diego Moores Cancer Center). Data acquisition was performed using Nikon NIS-Elements AR v5.21.01 software multi-color continuous mode. Single-molecule localization was achieved by rapid stochastic sequential laser (488 nm and 647 nm) activation and deactivation of a small subset of fluorophores at optimized power settings to enable localization of individual molecules. Sample drift was corrected using the software’s autocorrelation-based drift correction algorithm, and axial drift was minimized with the Nikon Perfect Focus System. Image acquisition was typically 1 to 2 million cycles of single molecule imaging and detection. Reconstruction of super resolution images, molecule localization visualization and data analysis were conducted using NIS-Elements N-STORM Analysis module. Sample preparation for STORM image acquisition was conducted according to the N-STORM Protocol-Sample Preparation guide and Dempsey et al. 2011 (46).

### Sucrose density gradient fractionation

Sucrose fractionation was carried out as previously described (10). EA.hy926 cells seeded at 4.95 x 10^6^ cells per 10-cm dish grown overnight, rinsed with ice cold PBS, and lysed in a sodium carbonate buffer (150 mM sodium carbonate at pH 11, 1 mM EDTA, and protease inhibitors). The lysates were processed by 10 strokes with a Dounce homogenizer, passed through an 18-gauge needle, and sonicated on ice (Branson Ultrasonics Corp). Cell lysates (800 µL) were mixed with an equal volume of 80% sucrose in MES-buffered saline supplemented with 300 mM sodium carbonate and placed in a 12-mL ultracentrifuge tube. Density gradients were created by adding 6 mL of 35% MES-buffered saline with 150 mM sodium carbonate on the top, and 4 mL of 5% sucrose in MES-buffered saline with 150 mM sodium carbonate on top of that. The samples were ultracentrifuged in an SW41 rotor for 20 h at 4°C with a force of 23,000 x g, and 1 mL fractions were collected sequentially, resolved by SDS-PAGE and immunoblotted as indicated quantified by densitometry using ImageJ software.

### Cell fractionation assay

HEK293 cells transfected with GRK5 WT, IFF, 4A or pcDNA3 plasmids only were grown on 15 cm dishes overnight. Cells were collected and suspended in detergent free hypotonic buffer (50 mM Tris-HCl, pH 8, 2.5 mM MgCl2, 1 mM EDTA) supplemented with protease inhibitors. Cells were sheared using a syringe with a 21-gauge needle ten times and cell lysates were centrifuged at 400 x g for 5 min. Supernatants were collected and subjected to ultracentrifugation at 150,000 x g for 20 min to isolate soluble/cytosolic fractions from particulate/membrane fractions as described (47). Equivalent amounts of soluble and particulate fractions were analyzed by immunoblotting and quantified by densitometry using ImageJ software.

### Bioluminescence resonance energy transfer assay

HEK293 GRK KO cells were seeded in a 6-well plate at a density of 5.0 x 10^5^ cells, grown overnight and transfected with PAR1-YFP (1000 ng), EPCR (500 ng), βarr2 Rluc (250 ng), and either GRK5 WT, IFF or 4A mutant, or pcDNA3.1 (125 ng). The following day, cells were reseeded into 96-well plates coated with poly-D-lysine at a concentration of 3 x 10^4^ cells per well and grown overnight. The cells were then washed with PBS, serum starved for 1 h using a 1:1 mixture of DMEM without phenol red and PBS. After the starvation, cells were preincubated with the RLuc substrate Coelenterazine H (5 μM) for 5 min and then stimulated with 20 nM APC. BRET measurements were taken using a Berthold TriStar LB941 multimode plate reader equipped with Micro WIN 2010 software using two filter settings: 480 nm for Rluc and 530 nm for YFP. The BRET signal was calculated as the emission at 530 nm divided by the emission at 480 nm. The BRET signals were then normalized to basal BRET ratios and expressed as a percentage over the basal level. The data were fit against a one-phase association nonlinear regression model, and the area under the curve (AUC) was calculated using Prism.

### Statistical Analysis and Software

Data were analyzed using Prism 10.4.1 (GraphPad Software) and Microsoft Excel software. Statistical analysis methods for data (mean ± S.D.) of three or more independent experiments are indicated in the figure legends, Student’s t-test was used to compare two groups, and one-way or two-way ANOVA was used for multiple comparisons. Figures were created in Adobe Illustrator and cartoons were created with BioRender.com.

## Acknowledgments

We thank all members of the Trejo laboratory for comments and advice. We especially thank Hannah Higa for Figure illustrations and Norton Cheng and Irina Kufarevav for constructive discussions and advice in experimental design.

## Author contributions

H.Q, L.O.C., C.B., M.A.L.-R., J.T conceptualization; H.Q., L.O.C., O.M.I. formal analysis, J.T. supervision; H.Q., L.O.C. validation; H.Q., L.O.C., M.G.R., O. M. I. investigation. H.Q., L.O.C. data curation; H.Q., L.O.C. visualization; H.Q., L.O.C., C. B., M.A.L.-R. and J. T. writing, review, and editing original draft; J.T. project administration, resources, and funding acquisition.

The authors declare no competing interest.

## Supporting information

There is no supporting information

## Abbreviations

The following abbreviations were used: APC, activated protein C; BRET, bioluminescence resonance energy transfer; Cav1, caveolin-1, EEA1, early endosomal antigen-1; EPCR, endothelial protein C receptor; CRISPR-Cas9, clustered regularly interspaced short palindromic repeats-CRISPR-associated protein 9; GPCR, G protein-coupled receptor; GRK5, GPCR kinase 5; IgG, immunoglobulin; PAR1, protease-activated receptor-1; Rluc, *Renilla* luciferase; Th, thrombin; YFP, yellow flourescence protein, WT, wildtype.

## Data Availability

The data are contained within the article.

